# Correction: Replication of BRIT1/MCPH1 G2/M Checkpoint Function Confirms Original Conclusions

**DOI:** 10.1101/2025.06.19.660578

**Authors:** Shiaw-Yih Lin, Hui Dai, Stephen J. Elledge

## Abstract

In our 2005 PNAS publication1, we reported that BRIT1/MCPH1 is required for proper activation of the G2/M checkpoint in response to DNA damage. A recent analysis using image comparison software, Proofig, identified a duplicated panel in Figure 2A, stemming from a figure assembly error in which data from a prior experiment involving a different gene (Claspin)2 were inadvertently reused. As the original data are no longer accessible due to the age of the experiment, we have independently repeated the G2/M checkpoint analysis. The new results fully confirm the original conclusion: BRIT1 depletion compromises G2/M arrest following ionizing radiation. This replication supports the integrity of our original findings and is provided in support of a formal correction requested by PNAS.

## Introduction

In the 2005 study published in PNAS^1^, we demonstrated that BRIT1 (also known as MCPH1), a gene implicated in primary microcephaly, plays a pivotal role in DNA damage checkpoint control. Specifically, we showed that BRIT1 is necessary for both intra-S and G2/M checkpoint responses following ionizing radiation (IR). A key piece of evidence supporting this conclusion was Figure 2A, which depicted FACS analysis of mitotic entry in U2OS cells following BRIT1 knockdown and irradiation.

Recently, a review of our data using Proofig software identified an inadvertent image duplication in Figure 2A. Upon investigation, we determined that an image from a prior Claspin-related experiment had been mistakenly used^2^. Given that the original data files are no longer available, we conducted an independent replication of the experiment to confirm the validity of our findings.

## Methods

U2OS cells were transfected with either control siRNA (MISSION® siRNA Universal Negative Control #1, SIC001) or esiRNA targeting BRIT1 (EHU047601, Millipore Sigma) using Lipofectamine 3000 (ThermoFisher Scientific). After 48 h, cells were either left untreated or exposed to 3 Gy of IR, followed by a 1-hour incubation.

Cells were fixed in 70% ethanol, permeabilized with 0.25% Triton X-100 in PBS, and stained using Alexa Fluor 647-conjugated phospho-histone H3 (Ser10) antibody (Cell Signaling Technology, 9716S). Following phospho-H3 staining, cells were incubated in propidium iodide solution to assess DNA content. Flow cytometry for DNA content and p-H3 staining was performed using a Beckman Coulter Gallios instrument at the MD Anderson Cancer Center Flow Cytometry and Cellular Imaging Core Facility, and data were analyzed with FlowJo v10. Protein knockdown was confirmed by Western blotting using BRIT1 (D38G5) Rabbit mAb from Cell Signaling Technology and ACTIN antibodies.

## Results

In agreement with our original findings, control siRNA-transfected cells exhibited robust suppression of mitotic entry upon irradiation (1.21% to 0.05% P-H3 positive cells). In contrast, BRIT1-depleted cells displayed impaired checkpoint activation, maintaining a higher mitotic fraction (1.15% to 0.42%). Western blot analysis confirmed efficient depletion of the BRIT1 protein.

These results replicate and reinforce the conclusion originally reported in PNAS: BRIT1 is essential for a functional G2/M checkpoint response to IR.

## Discussion

This replication was undertaken to correct an unintentional error in figure assembly from our 2005 PNAS publication. Although the original figure included a duplicated panel, our replicated experiment unequivocally confirms the original interpretation. Moreover, two independent studies have since confirmed BRIT1/MCPH1’s role in DNA damage-induced checkpoint regulation^3, 4^, further corroborating our conclusions.

We present these results here in accordance with PNAS editorial policy, which does not permit new data in formal corrections but allows linking to externally archived validation results.

---

**Legend.**
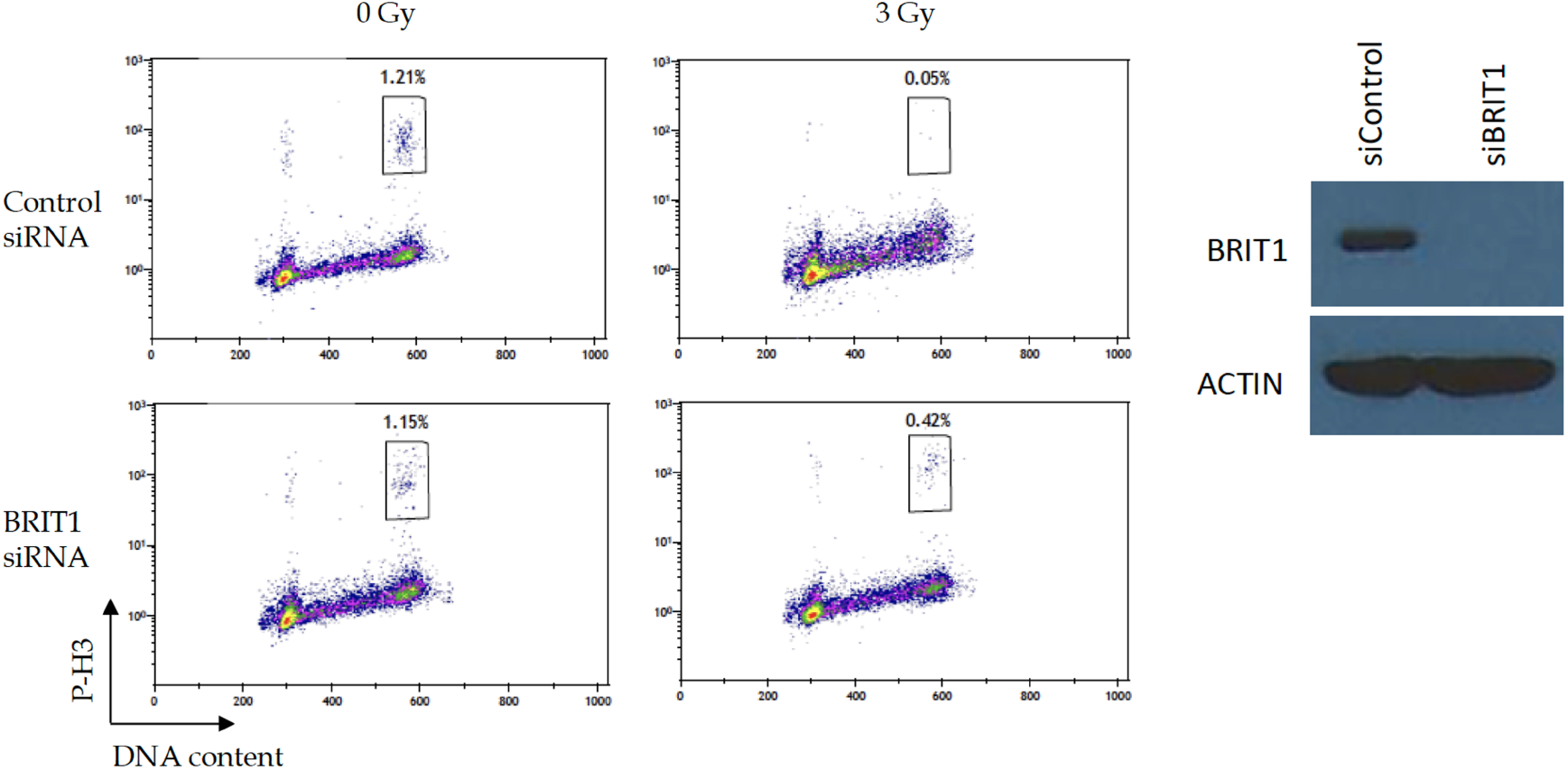
BRIT1 is required for the IR-induced G2/M checkpoint. (Left) U2OS cells were first transfected with either control or BRIT1 siRNA. Forty-eight hours after transfection, cells were either left untreated or irradiated with 3 Gy, followed by a 1-h incubation before fixation. Cells in mitosis were identified by staining with propidium iodide and a phospho-histone H3 antibody, followed by a FITC-conjugated secondary antibody. The percentage of M-phase cells was determined by FACS analysis for phospho-histone H3. (Right) Depletion efficiency was confirmed by western blot analysis with the indicated siRNAs.

